# Effects of the veterinary anthelmintic moxidectin on dung beetle survival and ecosystem functioning

**DOI:** 10.1101/213173

**Authors:** Paul Manning, Owen T. Lewis, Sarah A. Beynon

## Abstract

1. Macrocyclic lactones (MLs) are a class of chemical compounds administered to livestock for parasite control. These compounds are poorly metabolized and are predominately excreted in dung.
2. When coprophagous insects such as dung beetles (Coleoptera: Scarabaeoidea) are exposed to ML residues, lethal and sublethal effects are often observed. Indirectly this can lead to ML residues impairing ecosystem functions that underpin production. A strategy to reduce these negative effects involves selecting compounds that offer lower risk to non-target invertebrates such as the ML moxidectin.
3. Considering two dung beetle species with differing sensitivities to agricultural intensification, we asked whether exposure to moxidectin residues influenced survival, reproductive output, and functioning (short- and long-term estimates of dung removal).
4. When exposed to moxidectin, adults of the sensitive species (*Geotrupes spiniger* Marsham) experienced a 43% reduction in survival. In contrast, survival of the non-sensitive species (*Aphodius rufipes* L.) was unaffected. We were unable to determine whether exposure affected reproductive output of either species.
5. We found little evidence to suggest moxidectin impaired dung removal. We found however, that high densities of a species with relatively low functional importance (*A. rufipes*) can compensate for the loss of a functionally dominant species (*G. spiniger*). Over a longer timeframe, earthworms fully decomposed dung irrespective of moxidectin residues.

## Introduction

Macrocyclic lactones (MLs) are veterinary medicines (VMs) used to control internal and external parasites of livestock. These compounds function by agonizing chloride channels in the nervous tissue of invertebrates (Bloomquist, 1996) leading to hyperpolarization of nervous cells, flaccid paralysis, and death (Shoop & Soll, 2002). As MLs are highly effective, have negligible toxicity to mammals, and can be conveniently applied thorough a variety of formulations (Boxall *et al.*, 2007), they have become a widely-used tool for optimizing livestock health. Despite their popularity and efficacy, there is growing evidence that frequent use of MLs can have non-target environmental impacts on pasture-based production systems (Lumaret *et al.*, 2012). Macrocyclic lactones are poorly metabolized and active ingredients pass relatively unchanged into dung and urine (González Canga *et al.*, 2009). Invertebrates that colonize dung for food, breeding and shelter come into direct exposure with these residues. This can have both lethal (Beynon *et al.*, 2012) and sublethal (Verdú *et al.*, 2015) consequences for some taxa.

One such group is the dung beetles (Coleoptera: Scarabaeoidea), which play a key role in mediating dung decomposition (Nichols *et al.*, 2008). Death or impairment of dung beetles and other insects can slow rates of dung decomposition (Wall & Strong, 1987), which may lead to other detrimental consequences for ecosystem functioning in pastures. For instance, dung remaining unincorporated into the soil acts as a direct impediment to grass growth (Fincher, 1981). Delayed decomposition is also thought to impair the provision of numerous important ecosystem services (Beynon *et al.*, 2015), for example by increasing nitrogen loss through volatilization (Nichols *et al.*, 2008), increasing reinfection of livestock by parasites (Sands & Wall, 2016), and allowing pest flies to complete development (Bornemissza, 1970). Parasite resistance is less widespread for MLs compared to other classes of anthelmintic (Boxall *et al.*, 2003), so MLs are likely to remain a valuable tool for optimizing animal health. Finding ways to limit the non-target impacts of MLs on beneficial fauna and the functions they provide could contribute to improving the environmental and economic sustainability of livestock farming.

Limiting the non-target impacts of anthelmintics in the environment can be achieved through management decisions and mitigation measures, many of which are already practiced to slow parasite resistance to anthelmintics (Liebig *et al*., 2014). For instance, the selective treatment of livestock with high parasite burdens (van Wyk *et al.*, 2006) reduces exposure on a population level by providing sources of dung free from anthelmintic residues. Additional action can be taken under some management systems by housing animals during peak fecal excretion (Lumaret *et al.*, 2005). However, these management practices may be difficult to implement at a large scale. Identification of novel active ingredients and formulations with lower risks to non-target taxa remains an important goal for sustainable control of livestock parasites (Floate *et al.*, 2002; Beynon, 2012).

In this last regard, moxidectin is a ML that has shown promise. While moxidectin is effective in controlling a wide variety of livestock parasites (Losson & Lonneux, 1996), residues have greatly reduced impacts on non-target species compared to other MLs (Lumaret *et al.*, 2012). For instance, Doherty et al. (1994) found that moxidectin was required at 64 times greater concentration than abamectin (a related ML) to induce comparable LC_50_ mortalities in the dung beetle *Onthophagus taurus* (Schreber). The low toxicity of moxidectin relative to other MLs for non-target taxa has since been confirmed by numerous studies (e.g. Wardhaugh et al. 2001, Floate et al. 2005, Lumaret et al. 2012, Blanckenhorn et al. 2013). While moxidectin is a useful alternative to more toxic MLs, it would be misleading to assume that this compound poses negligible toxicological risk to all dung beetle species. Dung beetles exhibit significant interspecific differences in anthelmintic sensitivity (Beynon *et al.*, 2012), and failing to consider impacts on the more sensitive species could underestimate the non-target effects of anthelmintic treatment on structure and functioning of dung beetle communities. Although the mechanism is unknown, tunneling beetles tend to be more sensitive to anthelmintic residues than beetles from other functional groups (Beynon *et al.*, 2012). These dung beetles also play a key role in promoting dung decomposition (Roslin & Koivunen, 2001; Kaartinen *et al.*, 2013), leading to concern that the functions they support may be particularly susceptible to the effects of anthelmintic exposure.

Here, we investigate how moxidectin residues affect the functioning, survival, and reproductive output of two dung beetle species: a large-bodied tunneling species (*Geotrupes spiniger* Marsham) that has been shown to be sensitive to residues of an ML (ivermectin), and a smaller-bodied species (*Aphodius rufipes* L.) that is less sensitive to ivermectin (Beynon, 2012). These two species occur commonly during late summer throughout the Palearctic region.. The two species have different sensitivities to agricultural intensification (Hutton & Giller, 2003). To the best of our knowledge, their sensitivity to moxidectin has not previously been quantified or compared. We used manipulative experiments to investigate the influence of moxidectin residues on survival, reproductive output, and dung removal.

## Methods

The experiment was conducted on an improved pasture at Dr Beynon’s Bug Farm, St Davids, Pembrokeshire, UK (51°53′22″, 5°14′06″). The pasture was in long-term rotation, converted from arable (spring barley) production to pasture in 2006. The vegetation consisted of a mixture of perennial ryegrass (*Lolium perenne* L.) and white clover (*Trifolium repens* L.). At the time of the experiment, the field was grazed by a flock of 40 Welsh Mountain sheep and their lambs. Prior to beginning the experiment, we cut a 7×7 m area sward to a height of *c.*1cm. Sheep were excluded from the immediate experimental area with electric fencing.

Beetles were contained in 12L (N=36) enclosures, made from black plastic flower pots (Elixir Garden Supplies Ltd, Morecambe, Lancashire, UK) with the bases removed (Fig. 1). Each enclosure was dug into the soil to a depth of 8 cm which allowed colonization by soil invertebrates, which often play a key role in supporting dung removal (Kaartinen *et al.*, 2013). Enclosures were randomly assigned to one of three beetle treatments (*G. spiniger* added, *A. rufipes* added, or no beetles added). Each beetle treatment was replicated in two dung treatments (dung containing moxidectin residues, or dung from untreated cattle), giving a total of six treatments combinations with equal replication (n=6 replicates of each).

**Figure 1.**
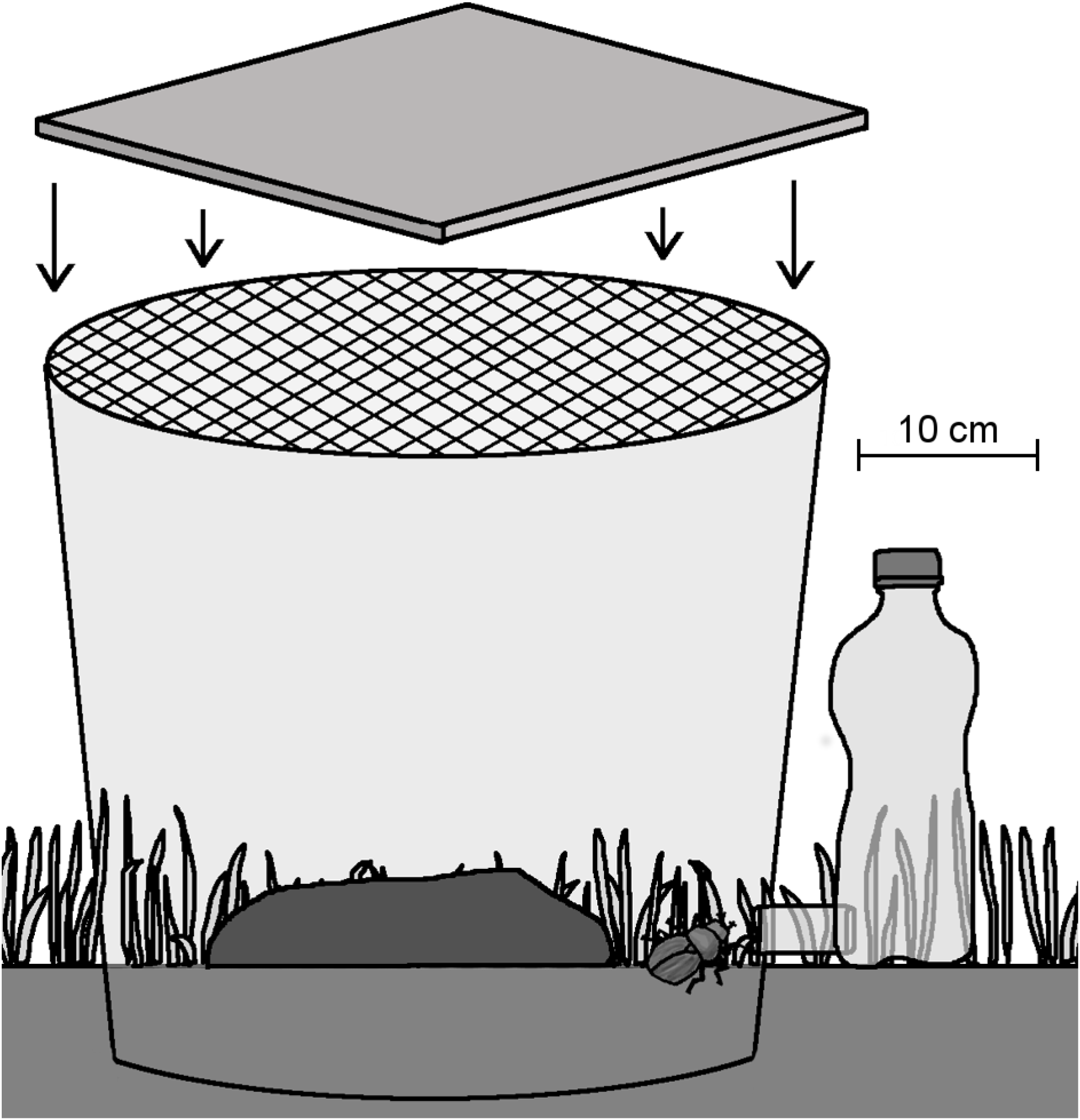
Depiction of the experimental enclosures. Soil invertebrates could freely colonize the enclosures from the underlying soil and contribute to dung removal.

Dung was collected from a group of Simmental (beef) cattle from a pasture adjacent to the experimental site (Upper Harglodd Farm, 51°53′23″, 5°14′07″). The cattle had not been treated with any VMs for the past four months. The experiment coincided with a scheduled treatment of a moxidectin product (Cydectin pour-on solution, 0.5% w/v moxidectin, Mole Valley Farmers, Camborne, Cornwall, UK) at the recommended dose of 1mL/10kg body weight. Fresh (<10 minutes old) dung was collected prior to anthelmintic treatment (control dung), and again nine and ten days following treatment (moxidectin dung), a period known to correspond to peak fecal concentration of moxidectin following pour-on application (Steel, 1998). Cattle grazed in the same field for the full duration of dung collection. All dung was frozen immediately after collection for a minimum of seven days to kill any insects that might have colonized it.

In the week leading up to the experiment, dung beetles were collected onsite through a combination of hand-searching cattle dung, and dung-baited pitfall trapping. Prior to the experiment, all beetles were housed in mixed-sex terraria containing cattle dung and sterilized potting soil, and kept in a well-ventilated shed.

The experiment began on August 23^rd^, 2014 when *G. spiniger* and *A. rufipes* were commonly observed flying onsite. Dung was thawed and homogenized, with control (Day 0) and treated (a mixture of 50:50 Days 9 and 10) dung kept separate. We measured 600g batches of dung with a kitchen balance accurate to 1g and, using a circular 12cm diameter plastic mold, formed dung pats using the methods of Beynon et al. (2012); a 6 cm space separated the dung from enclosure wall. Dung pats were placed on a 2cm aperture wire mesh, so that they could be lifted from the pasture surface, allowing non-destructive dung removal estimates in the field. Dung beetle biomass is a strong predictor of functioning (Nervo *et al.*, 2014), thus to allow more meaningful comparisons among treatments we standardized dung beetle biomass for all replicates using dry biomass (Table 1) following Beynon et al. (2012). Immediately after forming the dung pats, beetles were added to the enclosures and contained using a 2mm aperture plastic mesh.

**Table 1.** Dung beetle assemblages used within enclosures. Dung beetle dry biomass was approximately 1550mg in each treatment. Dry mass values are taken from Gittings and Giller (1997)^a^, and from Beynon (2012)^b^.

| Species | Functional group | Dry mass (mg) | Quantity | Total mass (mg) |
| --- | --- | --- | --- | --- |
| *Aphodius rufipes* (L.) | Soil-ovipositing endocoprid / kleptoparasite / paracoprid | 29.9<sup>a</sup> | 52 | 1555 |
| *Geotrupes spiniger* (Marsham) | Paracoprid | 387.0<sup>b</sup> | 4 | 1548 |

Enclosures (Fig. 1) were monitored four to five times per day for beetle activity. When beetles were first observed walking or flying within the enclosures (Day 14), we covered each enclosure with a ceramic tile to block light infiltration and fitted an emergence trap: a 500mL transparent plastic bottle attached using a 5cm length of 2cm diameter transparent plastic tubing. This encouraged beetles to move towards the light to exit the enclosure and enter the emergence trap. Enclosures were checked for emerging beetles three times a day for ten days (Days 14–24), until a four-day period without any additional emergence occurred (Days 20–24). At this point, we assumed no further beetles would emerge, removed the covers from all enclosures and replaced them with mesh coverings.

We estimated dung removal by measuring the remaining dung mass on Day 42, corresponding to a period when larval *A. rufipes* would have completed development within the dung pat and pupated in the soil (Stevenson & Dindal, 1985). As previous work has shown the benefits of beetles in supporting dung removal may become more apparent during the next grazing season (Beynon *et al.*, 2012; Nervo, B. *et al.*, 2017), we took a second measure of dung removal early in the next grazing season (May 14^th^, 2015). Enclosures were covered, and emergence traps were reattached on August 6^th^, 2015 to estimate impacts of moxidectin residues on larval development, as assessed by emergence of the F_1_ generation. Traps remained attached until September 30^th^, 2015, at which point we assumed that all adult beetles would have emerged.

### Analysis

The proportion of beetles that entered emergence traps was used to estimate beetle survival. The effect of moxidectin on adult survival was tested for each species by comparing the proportion emerged between the moxidectin residue treatment and the untreated-dung control by using a two-sample *t*-test. Dung removal was modelled as a function of dung beetle treatment (*G. spiniger*, *A. rufipes*, or beetles absent) and the presence or absence, of moxidectin using a two-way analysis of variance. Model residuals met assumptions of homogeneity and normality and thus raw values were left untransformed. Post-hoc comparisons amongst means were made using Tukey’s HSD. All analyses were carried out using R 3.1.1 (R Core Team, 2016). Figures were made using the ggplot2 package (Wickham, 2009).

## Results

Survival of *A. rufipes* was low in both dung types (control 0.66±0.06, treated 0.54 ± 0.07, mean ± se) and was not significantly affected by the presence of moxidectin residues (t = 1.29, df = 9.87, *P* = 0.23, Fig. 2). However, survival of *G. spiniger* was significantly affected by the presence of moxidectin residues (t = 4.77, df = 7.71, *P* = 0.002, Fig. 2), with survival reduced by approximately 43% in treated dung (0.54 ± 0.07) relative to the control (0.96 ± 0.04). No beetles from the F1 generation were successfully captured in emergence traps from either control or moxidectin residue dung.

**Figure 2.**
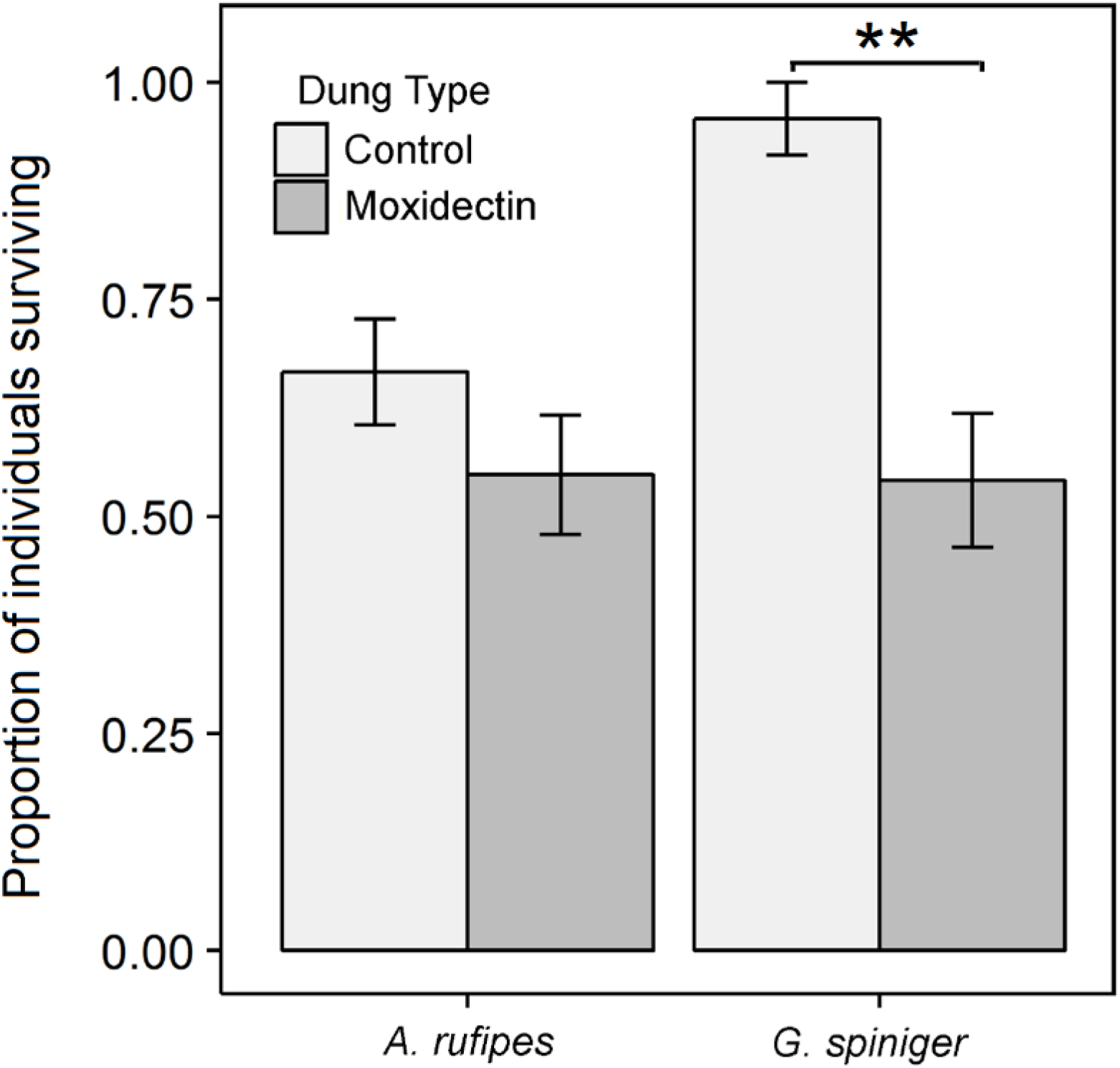
The proportion of surviving individuals captured in emergence traps, in response to moxidectin residues. *Aphodius rufipes* was not significantly affected. *Geotrupes spiniger* emergence was reduced by approximately 43% relative to controls. Bars are estimated means with standard errors.

For short-term dung removal, the interaction between dung type and beetle treatment was non-significant (F_2,30_ = 2.79, *P*=0.078). There was a significant main effect of the beetle treatment (F_2,32_ = 10.98, *P*<0.001) with significantly more dung removed when beetles were present compared to when beetles were absent. The presence of moxidectin residues did not significantly affect dung removal for either species (F_1,32_ = 0.37, *P* = 0.55). There was no significant difference in functional efficiency between the two species of dung beetle (Fig. 3).

**Figure 3.**
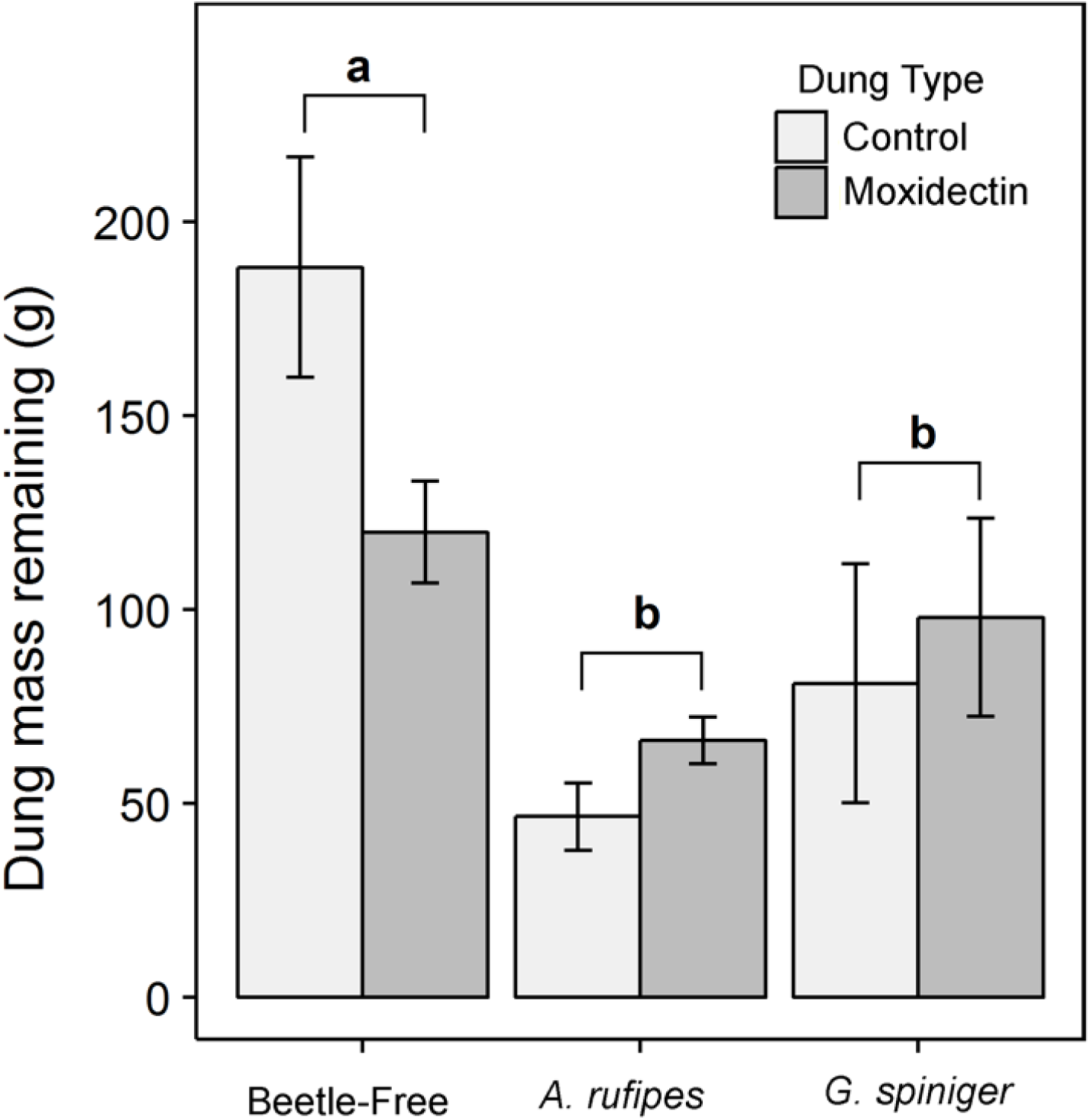
Mass of dung remaining (g) after six weeks as a function of the presence of dung beetles and moxidectin residues. Bars are estimated means with standard errors. Bars that do not share the same letter groupings are significantly different from one another.

It was not possible to estimate the effect that the beetle treatment and moxidectin residues had in explaining long-term functioning (dung removal at the beginning of the next grazing season), as dung was fully decomposed in all treatments, almost certainly due to earthworms: high densities of earthworm casts were observed within all enclosures.

## Discussion

Overall, we found little evidence that moxidectin impaired functioning both in the short and long term, but found that survival of one species (*Geotrupes spiniger*) was severely reduced by exposure to residues. In contrast to *G. spiniger, Aphodius rufipes* appeared to be unaffected by moxidectin residues (Fig. 2). This species tends to perform relatively poorly in enclosures, perhaps owing to its susceptibility to heat stress as it is a nocturnal species (Jessop, 1986).

In dung beetles, larvae are typically more sensitive to anthelmintic residues than adults (Beynon *et al.*, 2012). While we attempted to test how moxidectin residues affected larval development by quantifying emergence in the year following the experiment; we captured no beetles from the F1 generation. As we observed significant amounts of larval activity by *A. rufipes*, and evidence of brood ball partitioning by *G. spiniger* when estimating functioning in the short-term, it is unlikely that beetles failed to develop. It is more likely that our failure to recover any offspring in our experiment was a consequence of horizontal movement in the soil, with the subsequent generation emerging outside of the enclosures. It would be helpful to use enclosures dug deeper into the soil (e.g. Rosenlew and Roslin 2008), which limit horizontal movement in the soil, or larger enclosures to facilitate such movement, for future experiments. An additional explanation for the absence of emerging beetles in the second generation is the failure of beetles to develop within the timeframe of the experiment. While Palaearctic dung beetles are well-studied, detailed ecological and natural history information on the development of species in field conditions is scarce for most species (but see Landin 1961, Vessby 2001, Vessby and Wiktelius 2003, Floate et al. 2014). Understanding the environmental factors that limit the development of dung beetles within semi-field experiments such as ours would be useful for future work.

While we found no significant effect of moxidectin on short-term measures of dung removal (Fig. 2), this was likely due to high heterogeneity in the moisture content of dung, combined with a tendency for water content (as indicated by mass) to differ initially between the dung treatments. In the absence of beetles, the mass of dung containing moxidectin residues after 42 days (120 ± 13 g) was on average 37% lower than control dung (188 ± 28 g) although there was no significant difference attributable to the presence of moxidectin (t=-2.18, df =7.04, *P* = 0.065). A relatively small sample size, and variability in remaining dung mass (likely caused by the colonization of soil fauna) meant that we could not make any meaningful *post-hoc* corrections for differences in dung quality. More precise estimates of dung removal, including use of dry mass (Manning *et al.*, 2017), or loss-on-ignition (Menéndez *et al.*, 2016), would permit better tests of whether dung decomposition is impaired by anthelmintic residues, but either approach is destructive and would not allow us to make long-term estimates. Despite variability in initial dung conditions, we found that dung was fully degraded the following spring, regardless of the addition of beetles or the presence of moxidectin residues. This was likely due to the activity of earthworms, which can be highly effective in removing dung from pastures (O’Hea *et al.*, 2010), and have little sensitivity to moxidectin (Svendsen *et al.*, 2002), and other VMs (Kolar *et al.*, 2008).

In the short-term, we found that *A. rufipes* provided levels of dung removal equivalent to a similar total biomass of *G. spiniger*. Large tunneling dung beetles like *G. spiniger* are often considered to be the most important guild in supporting ecosystem functioning (Slade *et al.*, 2007; Kaartinen *et al.*, 2013; Nervo *et al.*, 2014). However, our results show that high densities of a small-bodied dung beetle (with similar total mass) can support equivalent levels of functioning in the short term. As differences in dung removal do not always correspond to differences in other beneficial functions in pasture (Manning *et al.*, 2016), studies that consider other relevant functions while maintaining dung beetle biomass constant, or using models that integrate body size (Nervo et al. 2014) might be useful in understanding how a high abundance of a small-bodied species could compensate for the absence of a large-bodied species often found to contribute disproportionately to functioning.

While dung beetles are important in supporting ecosystem functions that underpin pasture production, they represent only a small proportion of the diverse invertebrate assemblage associated with livestock dung in temperate agroecosystems (Skidmore, 1991) and are not the only taxa that contribute to functioning. Improved pasture in temperate climates can support dozens of earthworm (King *et al.*, 2008) and coprophagous fly species (Skidmore, 1991); both of which have been shown to make strong contributions to dung decomposition. For example, the earthworm *Eisenia fetida* (Savigny), the fly *Scathophaga stercoraria* L., and two species of *Aphodius* dung beetle larvae were all found to have similar functional importance in laboratory settings (O’Hea *et al.*, 2010). However, there are limited data on the relative importance of these taxa in supporting dung removal (Hendriksen, 1991; O’Hea *et al.*, 2010; Kaartinen *et al.*, 2013), and their roles in supporting associated ecosystem functions have not been widely explored. While economic models show that conserving dung beetles in agroecosystems may have considerable benefits to farmers (Losey & Vaughan, 2006; Beynon *et al.*, 2015), these estimates do not account for the role that other invertebrates play in supporting dung removal. If earthworms remove dung even in the presence of anthelmintic residues, high earthworm activity could compensate, in the longer-term, for functional deficits caused by the loss or impairment of more sensitive taxa.

Our findings suggest that while anthelmintic residues pose a threat to dung beetles in agroecosystems, death or impairment of these species may not always have marked functional consequences. For measures of short-term functioning (in the case of dung removal), high densities of a common species that is not sensitive to VMs may compensate for the absence of a more functionally important but sensitive species. Additionally, when the same function is considered over longer time periods, a combination of earthworm activity and abiotic weathering apparently compensates for the absence of dung beetles entirely. However, a more comprehensive assessment of functional impacts associated with treating livestock with moxidectin would need to consider multiple ecosystem functions and multi-species dung invertebrate communities. In addition, any persistent, multi-generational declines in dung beetles caused by repeated exposure should be investigated to fully appreciate the non-target risks of use of this anthelmintic in the livestock industry.

## Acknowledgements

We thank Mark and Emma Evans (Upper Harglodd Farm) for allowing us to collect dung for experiments. We also thank Andy Holcroft, Hannah Banks, and Sarah Sharpe for their assistance in the field. PM was funded by a Rhodes Scholarship, and a Natural Sciences and Engineering Research Council PGS (M) - Canada.

